# High-Quality Assembly of an Individual of Yoruban Descent

**DOI:** 10.1101/067447

**Authors:** Karyn Meltz Steinberg, Tina Graves Lindsay, Valerie A. Schneider, Mark J.P. Chaisson, Chad Tomlinson, John Huddleston, Patrick Minx, Milinn Kremitzki, Derek Albrecht, Vincent Magrini, Sean McGrath, Archana Raja, Carl Baker, Lana Harshman, LaDeana W. Hillier, Françoise Thibaud-Nissen, Nathan Bouk, Amy Ly, Chris Amemiya, Joyce Tang, Evan E. Eichler, Robert S. Fulton, Wesley C. Warren, Deanna M. Church, Richard K. Wilson

## Abstract

*De novo* assembly of human genomes is now a tractable effort due in part to advances in sequencing and mapping technologies. We use PacBio single-molecule, real-time (SMRT) sequencing and BioNano genomic maps to construct the first *de novo* assembly of NA19240, a Yoruban individual from Africa. This chromosome-scaffolded assembly of 3.08 Gb with a contig N50 of 7.25 Mb and a scaffold N50 of 78.6 Mb represents one of the most contiguous high-quality human genomes. We utilize a BAC library derived from NA19240 DNA and novel haplotype-resolving sequencing technologies and algorithms to characterize regions of complex genomic architecture that are normally lost due to compression to a linear haploid assembly. Our results demonstrate that multiple technologies are still necessary for complete genomic representation, particularly in regions of highly identical segmental duplications. Additionally, we show that diploid assembly has utility in improving the quality of *de novo* human genome assemblies.

## INTRODUCTION

High-throughput sequencing has enabled the characterization of thousands of human genomes in recent years; however, *de novo* high-quality assembly of many human genomes has remained difficult. This is due, in part, to the limitations of short reads that are unable to traverse repetitive elements longer than read length. Features such as segmental duplications, centromeres and telomeres are difficult to assemble and are also associated with gaps and errors in the reference genome assembly (Chaisson et al. 2015b). Yet, repeats and segmental duplications compose nearly half of the human genome, contain genes important to brain development (Dennis et al. 2012; Antonacci et al. 2014) and immune response (Watson et al. 2013, 2015), and are therefore vital to characterize in individual human genomes. Additionally, these loci often feature polymorphic copy number variants making it difficult to fully characterize this variation using reference-based methods that depend on a haploid consensus.

Technology advancements are making human *de novo* assembly a tractable problem. Long-read sequencing technologies generated from single molecules that are free from amplification steps have recently emerged as the frontrunners in advancing *de novo* assembly of complex genomes. For example, single-molecule, real-time (SMRT) sequencing from Pacific Biosciences (PacBio) generates reads from single molecules with read lengths greater than 10 kb. Mapping and local assembly of SMRT reads resolved or reduced almost 50% of the gaps in GRCh37 (Chaisson et al. 2015a). The IrysChip technology from BioNano Genomics (BioNano) linearizes DNA molecules of hundreds of kilobases, utilizes nicking enzymes to detect a seven-nucleotide sequence, and provides a genome map from this high resolution physical map.

Recently, Pendleton et al. (Pendleton et al. 2015) described a *de novo* assembly of a human genome of European ancestry using a mixture of PacBio and BioNano sequencing and genomic maps. From this hybrid assembly, they achieve a scaffold N50 of approximately 31 Mb and a contig N50 of 1.4 Mb with a total assembly size of 2.76 Gb. This is a nearly 2.5-fold improvement of scaffold N50 over other high-throughput sequence *de novo* assemblies (Gnerre et al. 2011) and a tenfold improvement of contig N50 over a recent Illumina-based *de novo* assembly of the single-haplotype hydatidiform mole CHM1 (Steinberg et al. 2014a). This hybrid approach was also used to assemble a genome from a Chinese individual with contig N50 of 8.3 Mb, scaffold N50 of 22 Mb, and total assembly size of 2.93 Gb (Shi et al. 2016). However, these assemblies are composed of sets of contigs that have not been scaffolded into chromosome assemblies and do not represent haplotype-resolved genome sequences.

10x Genomics (10x) also recently published on their partitioning technology that generates a novel data type known as “Linked-Reads” (Zheng et al. 2016). Briefly, high molecular weight genomic DNA is partitioned into 100,000 droplets, barcoded, and then released from the droplets. These molecules then undergo standard Illumina sequencing, and a downstream algorithm is able to parse the barcodes to link sequencing reads from the original DNA molecule resulting in linked reads with continuous stretches of phased variants. Mostovoy et al. recently described a method that combines Linked-Reads with BioNano map data to produce a diploid *de novo* assembly (Mostovoy et al. 2016). They achieve a scaffold N50 of 7.03 and a total genome size of 2.81 Gb of the NA12878 genome using this assembly technique. Recent improvements to the 10x technology, including increasing partitions to 4 million and improved biochemistry, allows for the production of a *de novo* diploid assembly from a single 10x library using the Supernova^™^ assembler (Jaffe et al., in preparation).

By applying the latest advances in sequencing and mapping technologies we can now accomplish *de novo* assembly of tens of individuals for the purpose of characterizing new variation not currently represented in the human reference assembly. One key advance has been the sequencing and finishing of single-haplotype human genomes (e.g., CHM1 and CHM13) (Steinberg et al. 2014a); (Schneider et al., in preparation). These data sets can be used to completely resolve complex genomic architecture and improve the GRC reference assembly. Additionally, the growth in clinical genome sequencing applications benefits from a human reference genome resource that accurately represents the diversity of the human population to facilitate the identification and characterization of disease-associated variants. The 1000 Genomes Project (1KG) (1000 Genomes Project Consortium et al. 2015) project provided a valuable foundation, yet it is now necessary to explore additional representative human genomes for deep sequencing, assembly, and finishing to high quality and contiguity.

In this paper we describe our efforts to create a *de novo* assembly of an African individual, NA19240, the daughter in a Yoruban trio sequenced as part of the 1KG Project, in the context of a larger effort to sequence and generate high-quality assemblies of human genomes from diverse populations to improve and add diversity to the reference assembly. We combine PacBio, Illumina, and BioNano technologies to create a highly contiguous assembly. We evaluate this assembly compared to the human reference assembly and other *de novo* assemblies. Finally, we explore how complementary technologies, such as BAC sequencing, Linked-Read data (Zheng et al. 2016) and novel algorithms that haploresolve assembly contigs (Chin et al. 2016), can be used to improve *de novo* assemblies.

## RESULTS

We sequenced NA19240 genomic DNA across 224 P6-C4 SMRT cells to generate approximately 49X coverage (147 Mb) with average read lengths of 12,331 base pairs. The data were error corrected using Quiver and assembled using FALCON (Chin et al. 2013), producing an assembly of 2.82 Gb with a contig N50 of 6,108,492 base pairs (see Table 1 for assembly metrics). Using stringent alignments of the NA19240 contigs to GRCh38, the assembly was ordered and oriented into chromosomes and submitted to GenBank as Hs-NA19240-1.0 (GCA_001524155.1). 97.9% of Hs-NA19240-1.0 was placed on chromosomes; the ungapped chromosomal length was 2.76 Gb, and the gapped chromosomal length was 3.03 Gb. There were 49,672,295 unplaced bases aggregated into unlocalized scaffolds for a total of 2.81 Gb of ungapped contigs.

**Table 1.** Assembly statistics

|  | # of Contigs | Contig N50 | Total Size (Gb) | # of Scaffolds | Scaffold N50 | Coverage |
| --- | --- | --- | --- | --- | --- | --- |
| NA19240 Falcon | 3569 | 6.01 Mb | 2.75 | NA | NA | 49X |
| BioNano | 3168 | 1.19 Mb | 2.85 | NA | NA | 72X |
| BioNano hybrid | NA | NA | 2.74 | 421 | 14.78 Mb | 72X |
| Hs-NA19240_1.0 | 3189 | 7.25 Mb | 3.08 | 1,588 | 78.6 Mb | 49X |
| DISCOVAR | 1,044,758 | 52,394 bp | 3.09 | 1,030,208 | 69.5 kb | 60.6X |
| DISCOVAR + SSPACE | 926,653 | 532,3 kb | 3.09 | 909,784 | 1.46 Mb | 181.9X* |
| NA19240_10x_dev** | NA | 100.5 kb | 2.73 | 1,530 | 16.09 Mb | 56X |

\*Sum of 60.6X (2x250bp OLR), 40X (3kb mate-pair), 81.3X (short-insert PE)
\*\*Statistics calculated using only scaffolds >10Kb

We then generated a 72X coverage genomic map using the BioNano Irys platform with molecules over 180kb in length for a total assembly size of 2.85Gb. The mapping rate of BioNano contigs to GRCh38 is 96.9% and is 95.9% to Hs-NA19240-1.0. After alignment of the BioNano map to the assembly, we identified inconsistencies using Illumina reads (see below) that could be the result of misassemblies or true variation and resolved them by breaking contigs. This process corrected 45 regions. We created a hybrid assembly (Materials and Methods) with the BioNano and FALCON assemblies that resulted in an assembly with a scaffold N50 to 14.78 Mb and total size of 3.74 Gb (Table 1).

We generated 42.4X PCR-free paired-end Illumina reads; 99.33% and 99.25% of reads can be aligned to GRCh38 and Hs-NA19240-1.0, respectively. We also generated 60.6X of 2x250 overlapping reads, 40X of 3 kb mate-pair, and 81.3X of short-insert (550 bp) paired-end Illumina data. We then performed *de novo* assembly of Illumina reads for the NA19240 assembly using the DISCOVAR *de novo* algorithm (Weisenfeld et al. 2014). This assembly had over one million contigs with a contig N50 of 52,394 bp and a total size of 3.09 Gb. The scaffolding and contiguity of the DISCOVARDeNovo assembly was then further improved by first running successive SSPACE (v. 3.0) (Boetzer et al. 2011) scaffolding jobs on the DISCOVARDeNovo scaffolds utilizing the additional 3kb mate-pair Illumina sequence. SEALER (Paulino et al. 2015) was used to enhance the scaffolding achieved from SSPACE as well as for merging of contig gaps.

Recently, a new approach to *de novo* assembly has been developed that produces haplotype-resolved assemblies by default. The Supernova™ assembler takes advantages of Linked-Reads to haplotype-resolve assemblies for an individual from a single library (Jaffe et al., submitted/in prep). An early assembly of the NA19240 individual (NA19240_10x_dev) from a development version of the Supernova v1.1 software was available and we wanted to assess whether the longer haplotype blocks, with an N50 of 11.41 Mb, available from this method improved gene annotation from the perspective of resolving frameshifting errors (Table 1).

### Assembly-Assembly and RefSeq Alignment

To evaluate the consensus quality of these assemblies, we aligned each assembly to one another using *nucmer* (Kurtz et al. 2004). Consensus accuracy between the GRCh38 primary assembly and the unscaffolded NA19240 assembly and Hs-NA19240-1.0 is 99.23% and 99.28%, respectively. The Hs-NA19240-1.0 and DISCOVAR+SSPACE NA19240 assembly has consensus accuracy of 99.67% while the DISCOVAR+SSPACE NA19240 assembly is 99.61% accurate when compared to GRCh38.

To further investigate the accuracy of the assembly, Hs-NA19240-1.0 was aligned to GRCh38.p2 using the NCBI pipeline for assembly-assembly alignment (Kitts et al. 2015). Briefly, NCBI alignments are generated in two phases. The first phase, or ‘First Pass’ alignments, are reciprocal best alignments, meaning any locus on the query assembly will have 0 or 1 alignment to the target assembly. ‘Second Pass’ alignments capture large regions (>1Kb) that have no alignment or conflicting alignments in the First Pass and represent duplicated sequences. In the ‘Second Pass’ alignments, a given region in the query assembly can align to more than one region in the target assembly, and these alignments can be used to identify regions of assembly collapse or expansion. Collapse indicates fewer copies of a sequence, while expansion suggests more copies of a sequence. These events can occur either due to assembly error or to biological differences between the two samples contributing to the assemblies. The First Pass alignments indicate that 98.6% of Hs-NA19240-1.0 is covered by reciprocal best alignments to GRCh38.p2, while 88.4% of GRCh38.p2 is covered by reciprocal best alignments to Hs-NA19240-1.0, and that the overall average percent identity is 99.73% between the two assemblies (based only on first pass). There are approximately 4.2Mb of gapped bases. This overall identity is slightly higher than the *nucmer* consensus accuracy due to slight differences in alignment methods.

Examining the percent of Second Pass alignments where each alignment is counted suggests that there is more collapse in Hs-NA19240-1.0 when compared the GRCh38 reference. Across all primary chromosomes, there is an average of 3.57% of Second Pass alignments consistent with the propensity of whole-genome sequence (WGS) assemblies to exhibit collapse of highly related sequences, compared to clone-based assemblies. For comparison, we also plotted the percent of Second Pass alignments from the assembly-assembly alignment of CHM1_1.1 to GRCh38.p2 (Figure 1). This WGS assembly has a lower average percent of second pass alignments (1.49%) compared to Hs-NA19240-1.0 due to the resolution of large regions of segmental duplication in CHM1_1.1 with high-quality BAC sequence tiling paths (Steinberg et al., 2014a). These results together suggest that although the PacBio assembly is highly contiguous compared to short-read assemblies, there are still regions that cannot be resolved without BAC sequencing (see Large-Insert Clone Analysis section below).

**Figure 1.**
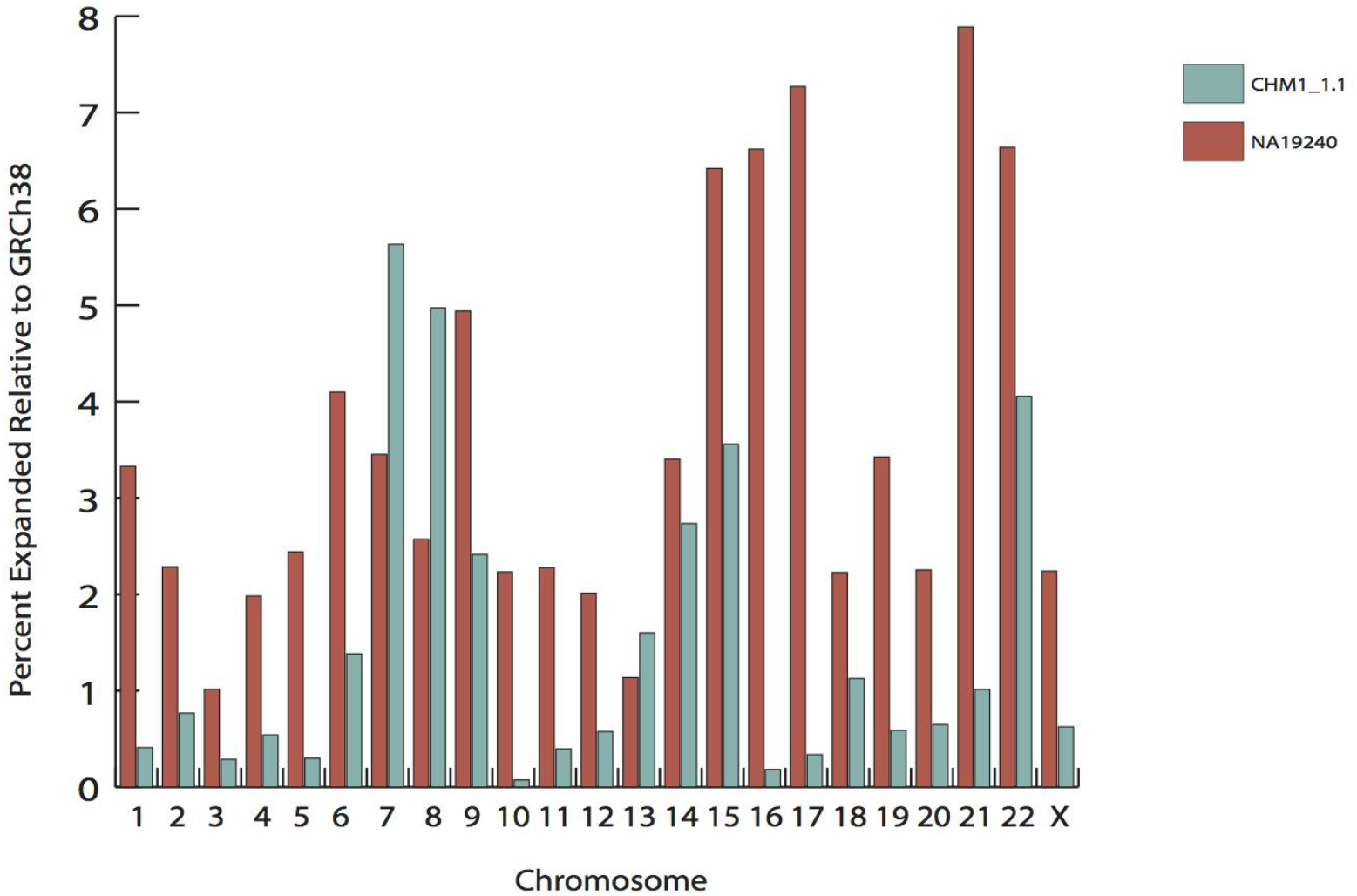
Second Pass alignments of Hs-NA19240-1.0 and CHM1_1.1 to GRCh38. Second pass alignments show that Hs-NA19240-1.0 has more collapse than CHM1_1.1 relative to GRCh38.p2 consistent with other WGS assemblies that do not resolve segmental duplications with BAC sequences.

We then aligned a set of 56,130 human RefSeq transcripts to GRCh38, Hs-NA19240-1.0, and NA19240_10x_dev. The native output of the Supernova^™^ assembler is a graph structure (with a scaffold N50 of 16.9Mb) that must be converted to FASTA for input into most pipelines. To create a haploid FASTA representation, a process traverses the graph and selects one haplotype at each haplotype block (‘megabubble’, http://support.10xgenomics.com/de-novo-assembly/software/pipelines/latest/output/generating). We generated two independent sets of haploid FASTA files to represent the diploid assembly. As the assessment pipeline is not prepared to deal with diploid assemblies, we aligned each haplotype separately as an independent assembly (hap2.1 and hap2.2) (Table 2). Overall, >98% of sequences aligned to the Hs-NA19240-1.0 with only 0.2% of sequences represented as multiple alignments or split transcripts. The percentage of transcripts rejected because they are co-placed with transcripts representing different genes is a way to measure assembly collapse and is between 0.41-0.56%. This suggests that Hs-NA19240-1.0 has collapsed sequences from paralogous gene copies and is consistent with the second pass alignment analysis. This is consistent with other PacBio-based assemblies that have been generated as part of a larger effort to generate high-quality human reference assemblies (Schneider et al., in preparation).

**Table 2.** RefSeq alignment of two different NA19240 assemblies

| Name | GRCh38 | Hs_NA19240-1.0 | hap2.1 | hap2.2 |
| --- | --- | --- | --- | --- |
| Accession | GCF_000001405.26 | GCA_001524155.1 | NA | NA |
| Number of sequences retrieved from Entrez | 56130 | 56130 | 56130 | 56130 |
| Number align-able sequences | 56130 | 55832 | 55832 | 55832 |
| Number of sequences not aligning | 26 | 608 | 366 | 363 |
| Number of coding transcripts retrieved from Entrez | 43217 | 43217 | 43217 | 43217 |
| Number align-able coding sequences | 43217 | 43068 | 43068 | 43068 |
| Number of rank 1 aligning coding transcripts | 43182 | 42589 | 42832 | 42833 |
| Number of coding sequences not aligning | 21 | 414 | 134 | 134 |
| Number of noncoding transcripts retrieved from Entrez | 12913 | 12913 | 12913 | 12913 |
| Number align-able noncoding sequences | 12913 | 12764 | 12764 | 12764 |
| Number of rank 1 aligning noncoding transcripts | 12903 | 12499 | 12500 | 12503 |
| Number of noncoding sequences not aligning | 5 | 194 | 232 | 229 |
| Number (%) of sequences with multiple best alignments (split transcripts) | 13 (0.02) | 99 (0.18) | 1914 (3.46) | 1891 (3.42) |
| Number (%) of sequences with CDS coverage < 95% | 21 (0.05) | 1061 (2.49) | 2260 (5.28) | 2261 (5.28) |
| Number of coding transcripts dropped at consolidation | 0 | 327 | 401 | 401 |
| Number of noncoding transcripts dropped at consolidation | 0 | 232 | 287 | 283 |
| Number of annotated proteins with frameshifts | 19 | 8225 | 225 | 218 |
| Number of annotated proteins with non-frameshifting indels | 18 | 490 | 489 | 508 |

We examined genes with indels and categorized them as not frameshift (NFS; those having indels that are multiples of three nucleotides that would not disrupt frame) or true frameshifts (Table S1). Frameshifts can be an indication of over-scaffolding in an assembly (Florea et al. 2011). The GRCh38 primary assembly has 19 frameshifted transcripts. There are 225 and 218 frameshifted transcripts in hap2.1 and hap2.2, respectively, but only 120 transcripts frameshifted on both haplotypes. Of these 120 transcripts, 66 were also frameshifted in the Hs_NA19240-1.0 assembly, suggesting a possible biological basis for this observation. There are 48 transcripts frameshifted in Hs-NA19240-1.0 and hap2.1 but not hap2.2, and there are 112 frameshifted transcripts in Hs-NA19240-1.0 and hap2.2 but not hap2.1. Eight of the 52 unique genes (from 66 transcripts) that are frameshifted in both assemblies overlap segmental duplications. Hs_NA19240-1.0 has the most frameshifted transcripts (8,225 transcripts from 4,132 unique genes). Most genes were only frameshifted in one assembly (n=4,071), but there were two genes (three transcripts) frameshifted in all four assemblies: *ABO* and *PKD1L2*, the latter a known processed pseudogene. Although there is an excess of frameshifted transcripts in Hs-NA19240-1.0 compared to hap2.1 and hap2.2 from NA19240_10x_dev, the numbers of transcripts (both coding and noncoding) that align to each assembly are similar (Table 2). In addition, the numbers of transcripts dropped at consolidation are similar between the three assemblies. There are only 17 transcripts present in Hs-NA19240-1.0 that are dropped in hap2.1 and hap2.2 suggesting collapse in a similar fashion and that the genomic content between Hs-NA19240-1.0 and NA19240_10x_dev is comparable. The excess of frameshifts in Hs-NA19240-1.0 may be due to the linear compression to a haploid consensus that creates mosaic alleles that are not representative of the diploid state. The significantly lower numbers of frameshifted transcripts in NA19240_10x_dev (both haplotypes) are suggestive of this explanation and may also be due to the higher read accuracy of Illumina sequencing technology.

### Repetitive Element Content

We ran RepeatMasker (Smit et al. 1996) on both the primary assembly of sequence that was placed on chromosomes and the primary assembly plus the unplaced scaffolds of the HS-NA19240-1.0 assembly. 44.53% of the primary assembly is masked with interspersed repeats (the largest categories being L1 LINEs at 16.46% and Alu elements at 9.68%) while an additional 1.67% of the primary assembly is composed of low complexity, satellite and simple repeats (Table S2). Adding the unplaced scaffolds only increased repetitive content by 0.24%; 100% of the additional repetitive content was alpha-centromeric satellite.

Segmental duplications were assessed using Whole Genome Assembly Comparisons (WGAC) (She et al. 2004) and Whole Genome Shotgun Sequence Detection (WSSD) (Bailey et al. 2002). Nearly 5% of the genome is predicted as duplicated using both methods. The total amount of nonredundant duplication sequence identified by WGAC is 112.84 Mb, and WSSD identifies 74.48 Mb of duplicated sequence. Sixty-five percent of WSSD duplications are shared with WGAC duplications greater than 10kb and greater than 94% identity. Approximately 40% of segmental duplications in Hs-NA19240-1.0 are present in the unplaced scaffolds and represent highly identical tandem duplications (Figure S1).

### Large-Insert Clone Analysis

#### Fosmid library

Fosmid clones from the ABC10 library generated from NA19240 DNA (Kidd et al., 2008) were aligned to the NA19240 assembly as well as to GRCh38 using the NCBI end alignment pipeline (Schneider et al. 2013a). Clones are placed if both ends of the clone hit a common contig. Concordant clones have both ends correctly oriented and a calculated placement length within three times the standard deviation of the mean clone size. Clones are placed either uniquely or multiple times in an assembly unit. 723,116 clones were placed on Hs-NA19240-1.0 representing 71.6% of clones that had both ends assembled (1,008,980). Of those, 719,480 are uniquely placed; 97.6% of the uniquely placed clones are concordant. In comparison, 96% of CH17 clone placements on CHM1_1.1 are unique and concordant. These data indicate the high quality of these assemblies, as we expect contiguous assemblies to place 95% or greater of clones as unique and concordant. 739,519 ABC10 clones were placed on GRCh38 representing 73.3% of total clones with both ends. There are approximately twice the number of clones with multiple placements on GRCh38 (7,632) compared to Hs-NA19240-1.0 (3,636). The clone inserts with multiple placements on GRCh38 also have a twofold enrichment of segmental duplication content compared to Hs-NA19240-1.0 likely due to the unresolved nature of segmental duplications in Hs-NA19240-1.0 (see above). This is also reflected in the fact that there are more discordant clones placed on Hs-NA19240-1.0 (17,516) compared to GRCh38 (14,549) suggesting unresolved structural variation in the assembly.

#### BAC Library

We created a BAC library (VMRC64) from NA19240 DNA. We identified 784 BAC clones from five regions of known structural variation and/or gaps in the reference assembly and selected 93 clones for sequencing (Table S3). In addition, we created 96 probes for 16 regions of human-specific duplications; we identified 1,209 total clones and selected 76 for sequencing (Table S3). We will be integrating these clone paths into the next iteration of the NA19240 assembly when they are finished to improve contiguity and resolve complex genetic architecture. For example, there is a gap in Hs-NA19240-1.0 at the *NBPF11* locus on chromosome 1 due to highly identical segmental duplications in this region (Figure 2).

**Figure 2.**
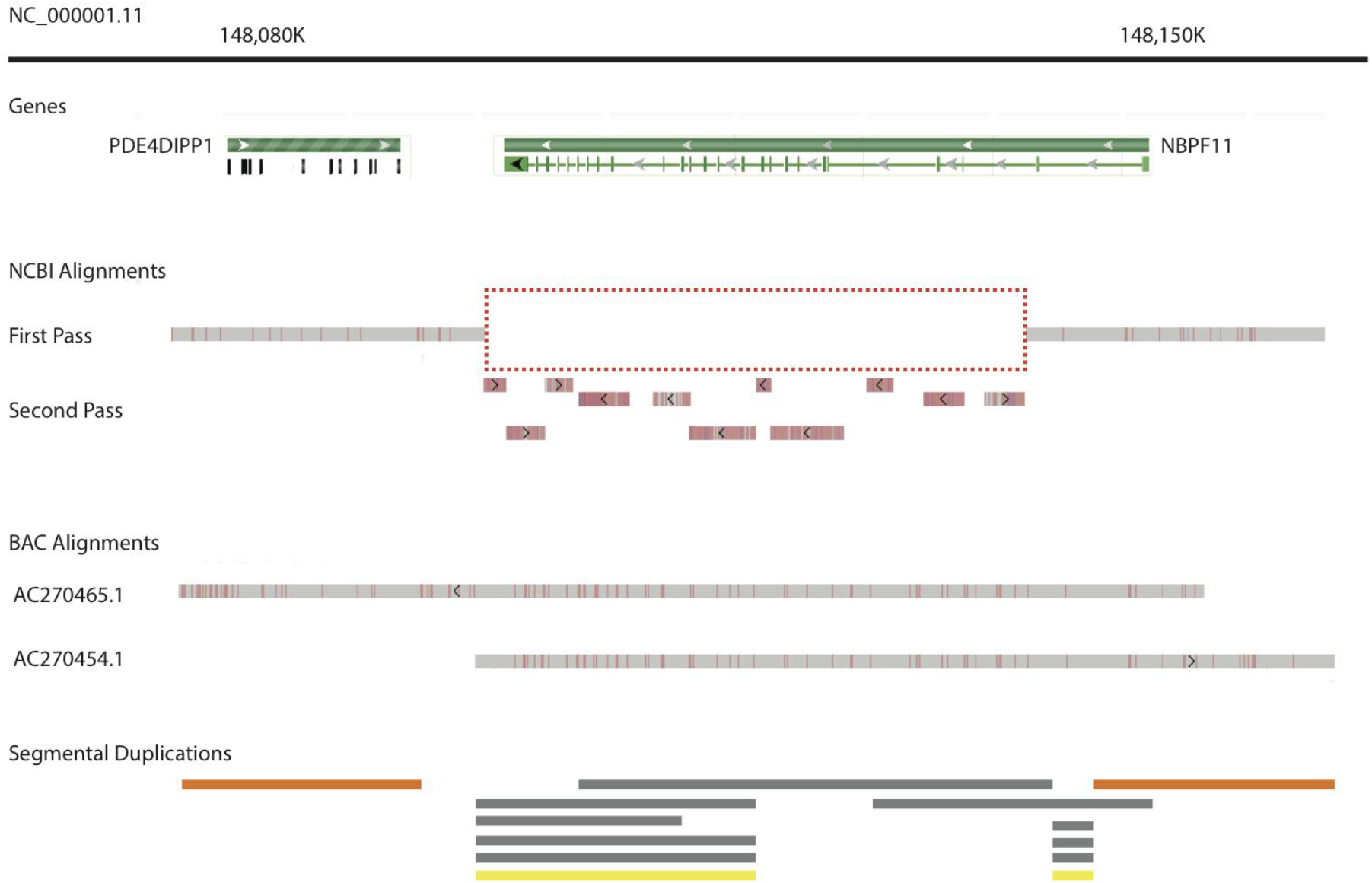
Using BACs to fill gaps and fix misassemblies. The NBPF11 region of NC_000001.11 on GRCh38.p2 is shown (chr1:148,080,000-149,150,000) with the NCBI first and second pass alignments of NA19240 contigs, BAC alignments and segmental duplications. The first pass alignment shows a gap over most of *NBPF11* (highlighted in red dotted box) while the second pass alignments show many small alignments to various NA19240 contigs that are a result of significant segmental duplication content in this region. Two BAC clones, AC270465.1 and AC270454, form a tiling path that covers this region. *NBPF11* is now completely represented in the Hs-NA19240-1.0.

Two BAC clones, AC270465.1 and AC270453.1, create a single path across this region and fill the gap from the WGS assembly. In another example, a clone path (AC270311 and AC270312) was aligned to Hs-NA19240-1.0 at the beta defensin *(DEFB)* locus on chromosome 8 where the WGS assembly was fragmented. This locus is known to harbor extreme structural variation and is not properly assembled in the reference assembly. The BAC clones aligned in reverse orientation to two regions of the Hs-NA19240-1.0 assembly suggesting that there was an inversion of these contigs (Figure S2). This inversion may represent true biology as NA19240 is homozygous for a 4.2Mb inversion of this locus while the reference assembly is in direct orientation (Antonacci et al. 2009). Additionally, the misassembly in the reference may also be contributing to this structural variant as well since Hs-NA19240-1.0 is aligned to the reference during scaffolding to chromosomes. It will take manual review of each WGS contig and BAC alignment to determine the correct tiling path through this region, and we will incorporate this path into the next iteration of Hs-NA19240-1.0. At this time, we have completed one path of structurally variant/gap regions and nine paths of human-specific duplications.

### Structural Variation

BioNano data generated from NA19240 DNA material was aligned to GRCh38, Hs-NA19240-1.0, and NA19240_10x_dev (hap2.1 and hap2.2) and structural variants (SVs) were called. There were 1,007 SVs called against GRCh38 and 1,238 SVs called against Hs-NA19240-1.0. Approximately 17% of the GRCh38 SVs were filtered and 23% of the NA19240 SVs were filtered from analysis to exclude duplicate SV calls and complex or small (<1kb) SVs. This resulted in 957 SVs called against GRCh38 (188 >10kb) and 840 SVs called against Hs-NA19240-1.0 (148 >10kb) (Table 3). The average size of GRCh38 insertions and deletions is 32,997 bp, the average size of Hs-NA19240-1.0 insertions and deletions is 70,379 bp. There were significantly more SVs called against NA19240_10x_dev, particularly as ends, insertions, and insertions of “N” bases due to inaccurate gap sizing in the 10x assembly software. Ends are called when a BioNano map extends beyond the end of the alignment from a contig/scaffold where the extending tail does not align anywhere else in the genome. There are more translocations called on Hs-NA19240-1.0 compared to the GRCh38 (46 vs 24) assembly. Translocations are called when a BioNano map extends beyond the end of the alignment from a contig/scaffold and the extending tail has an alignment somewhere else in the genome. It may indicate a chimeric scaffold, a potential join between two scaffolds or may be hitting a segmental duplication. There are 15 intrachromosomal and 31 interchromosomal translocations called against Hs-NA19240-1.0. Of the 31 interchromosomal events, 12 are translocations between a chromosome on the primary and an unplaced scaffold. All 46 of the translocations overlap segmental duplications and suggest that these loci are misassembled in Hs-NA19240-1.0.

**Table 3.** Filtered BioNano structural variants

|  | GRCh38 | Hs-NA19240-1.0 | hap2.1 | hap2.2 |
| --- | --- | --- | --- | --- |
| deletion | 237 | 82 | 27 | 28 |
| deletion_nbase | 24 | 123 | 63 | 60 |
| end | 191 | 272 | 744 | 729 |
| insertion | 421 | 114 | 309 | 317 |
| insertion_nbase | 2 | 160 | 1957 | 1958 |
| inversion | 20 | 6 | 1 | 1 |
| inversion_nbase | 10 | 19 | 7 | 7 |
| inversion_partial | 32 | 25 | 8 | 8 |
| translocation_interchr | 13 | 25 | 212 | 213 |
| translocation_intrachr | 7 | 14 | 0 | 0 |
| TOTAL | 957 | 840 | 3328 | 3321 |

Overall, there are fewer indels called against Hs-NA19240-1.0 than the GRCh38 assembly as expected, and indels called on Hs-NA19240-1.0 may represent variation missed by scaffolding our assembly using GRCh38. For example, at the *KIF5C* locus, NA19240 more closely matches the alternate scaffold with a 6 kb insertion (NW_003571033.2) than the primary reference (GRCh38, chr2:148K-150K). This is a known SV (GRC Issue HG-1068) where a fosmid clone, AC206420.3 (ABC10-48936800O12), has a 6,087 bp insertion relative to the reference.

However, due to linear compression, the non-inserted allele is represented in Hs-NA19240-1.0, and the insertion is called in this analysis.

Second, we used Assemblytics (Nattestad and Schatz 2016) to detect SVs directly from *nucmer* alignments of homologous chromosome scaffolds of GRCh38 (alt plus decoy) and Hs-NA19240-1.0. We identified 12,029 SV events (indels, tandem and repeat expansions and contractions >50bp) that affected a total of 7,156,888 bp (Table 4). There was a predictable enrichment for ~300 bp indels that aligned to Alu sequences and for ~6kb indels that aligned to LINE elements.

**Table 4.** Assemblytics results

|  |  | Count | Total bp |
| --- | --- | --- | --- |
| Insertion | 5-10 bp | 39565 | 257236 |
|  | 10-50 bp | 28894 | 493016 |
|  | 50-100 bp | 1064 | 72859 |
|  | 100-1,000 bp | 1707 | 472143 |
|  | 1,000-10,000 bp | 352 | 1063160 |
|  | Total: | 71582 | 2358414 |
| Deletion | 5-10 bp | 49582 | 320615 |
|  | 10-50 bp | 35029 | 609740 |
|  | 50-100 bp | 1150 | 76044 |
|  | 100-1,000 bp | 1707 | 488933 |
|  | 1,000-10,000 bp | 318 | 1104845 |
|  | Total: | 87786 | 2600177 |
| Tandem Expansion | 5-10 bp | 59 | 392 |
|  | 10-50 bp | 134 | 3980 |
|  | 50-100 bp | 281 | 20635 |
|  | 100-1,000 bp | 1935 | 734550 |
|  | 1,000-10,000 bp | 430 | 980970 |
|  | Total: | 2839 | 1740527 |
| Tandem Contraction | 5-10 bp | 29 | 198 |
|  | 10-50 bp | 147 | 5419 |
|  | 50-100 bp | 421 | 30009 |
|  | 100-1,000 bp | 1049 | 294299 |
|  | 1,000-10,000 bp | 143 | 372718 |
|  | Total: | 1789 | 702643 |
| Repeat Expansion | 5-10 bp | 94 | 595 |
|  | 10-50 bp | 103 | 2692 |
|  | 50-100 bp | 84 | 6278 |
|  | 100-1,000 bp | 531 | 214952 |
|  | 1,000-10,000 bp | 278 | 778623 |
|  | Total: | 1090 | 1003140 |
| Repeat Contraction | 5-10 bp | 10 | 68 |
|  | 10-50 bp | 78 | 2325 |
|  | 50-100 bp | 101 | 7656 |
|  | 100-1,000 bp | 624 | 209330 |
|  | 1,000-10,000 bp | 159 | 518129 |
|  | Total: | 972 | 737508 |

Third, we applied a modified version of SMRT-SV (Chaisson et al. 2015a)(Huddleston et al., in preparation) that detects SV signatures and develops local assemblies. We identified 12,728 insertions and 9,829 deletions that affected a total of 10,562,662 bp (SV by chromosome in Figure S3). The large majority of repeat-associated SVs are associated with tandem repeats (TRs; 61% of insertions and 46% of deletions; Table 5). Many insertions and deletions were not associated with repetitive elements (19.6% of insertions and 24.2% of deletions).

**Table 5.** Structural variant calls from SMRT-SV sorted by repeat element

| Class | Insertion |  |  |  | Deletion |  |  |  |
| --- | --- | --- | --- | --- | --- | --- | --- | --- |
|  | Count | Average | Std. dev. | Total | Count | Average | Std. dev. | Total |
| Not repetitive | 2489 | 616.96 | 1896.56 | 1535611 | 2382 | 617.23 | 1706.86 | 1470250 |
| Complex | 391 | 2252.44 | 2626.86 | 880705 | 383 | 2909.28 | 3551.8 | 1114255 |
| AluY | 825 | 288.22 | 62.45 | 237785 | 1248 | 283.74 | 79 | 354107 |
| AluS | 65 | 183.15 | 82.66 | 11905 | 116 | 153.33 | 92.67 | 17786 |
| L1Hs | 147 | 3160.68 | 2354.79 | 464620 | 143 | 2962.95 | 2531.6 | 423702 |
| L1 | 96 | 926.25 | 1538.46 | 88920 | 113 | 731.93 | 1312.18 | 82708 |
| L1P | 100 | 979.12 | 1405.82 | 97912 | 136 | 888.58 | 1323.03 | 120847 |
| SVA | 207 | 770.54 | 1049.79 | 159501 | 169 | 808.64 | 1046.73 | 136661 |
| HERV | 28 | 492.61 | 764.53 | 13793 | 37 | 464 | 804.22 | 17168 |
| Alu-mosaic | 130 | 427.06 | 251.51 | 55518 | 96 | 288.46 | 228.12 | 27692 |
| STR | 259 | 152.2 | 222.44 | 39419 | 138 | 92.3 | 79.87 | 12738 |
| Satellite | 88 | 275.86 | 376.38 | 24276 | 130 | 185.59 | 261.18 | 24127 |
| Singletons | 138 | 167.7 | 309.44 | 23143 | 177 | 122.37 | 155.47 | 21659 |
| Tandem_Repeats | 7707 | 291.92 | 507.59 | 2249822 | 4515 | 180.06 | 279.88 | 812951 |
| Total | 12728 | 464.35 | 1186.58 | 5910194 | 9829 | 473.34 | 1333.27 | 4652468 |

We then compared the SV calls from all three callers (Figure 3). Although BioNano did not call as many SVs as SMRT-SV, 87.7% of the SV calls intersected SMRT-SV calls. BioNano does not perform well on small SVs (<1.5kb), while SMRT-SV calls SVs down to 50bp, likely explaining the discrepancy. 48.2% of SV calls from Assemblytics intersect with SMRT-SV, but only 21.3% of SMRT-SV calls intersect with Assemblytics calls. Additionally, roughly two-thirds of BioNano calls intersect Assemblytics, but less than 5% of Assemblytics calls intersect BioNano calls.

**Figure 3.**
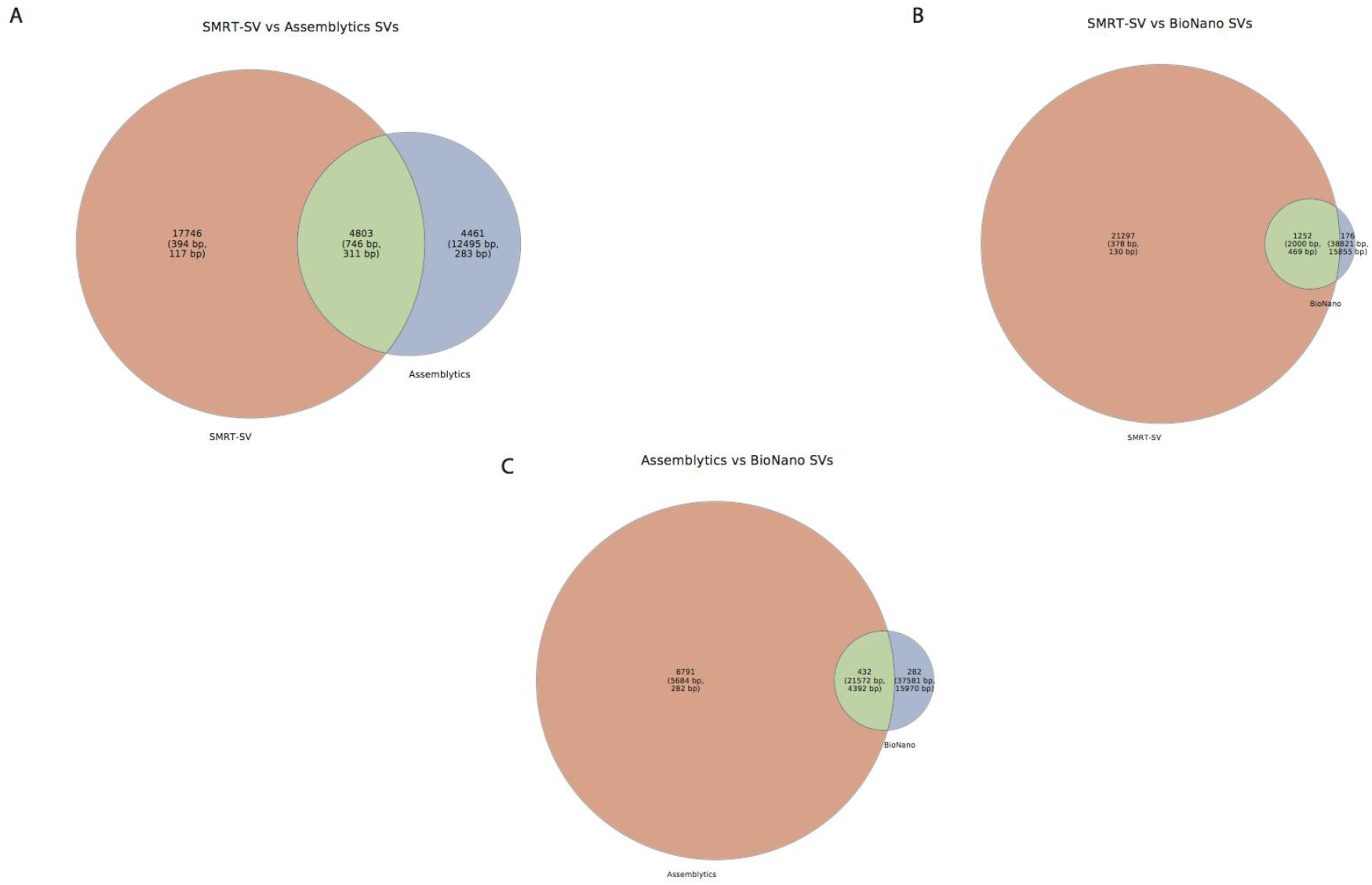
Comparison between BioNano, Assemblytics, and SMRT-SV. Venn diagrams were generated from these intersections and show the number of calls shared between the two callers as well as the mean and median SV size for each Venn panel. (A) shows the intersection between SMRT-SV and Assemblytics, (B) shows the intersection between SMRT-SV and BioNano, and (C) shows the intersection between Assemblytics and BioNano SVs.

When we compared Assemblytics and SMRT-SV deletions, we found that 72% of all Assemblytics deletions are in TRs, 85% of Assemblytics-only deletions are in TRs and 61% of Assemblytics deletions shared with SMRT-SV are in TRs. 50% of Assemblytics-only deletions are within 1kb of SMRT-SV deletion calls and 94% of those are in TRs. The pattern for insertions was almost identical with 59% of Assemblytics-only insertions within 1kb of a SMRT-SV insertion with 84% in TRs. However, if adjacent calls were required to have no more than 100bp difference in size, only 27% of Assemblytics-only deletions and 15% of Assemblytics-only insertions match those constraints. Based on these results, it is likely that variability in SV calling in TRs are responsible for a substantial proportion of Assemblytics-only events, although most of those Assemblytics-only events are different calls from adjacent SMRT-SV events.

### Resolving haplotypes with FALCON-Unzip

We used the haplotype resolving tool, FALCON-Unzip on Hs-NA19240-1.0 to represent the diploid genome as correctly phased haplotype contigs, or “haplotigs.” (Chin et al. 2016). The string graph from FALCON contains haplotype-fused contigs (no major SV) and bubbles (major SV). Briefly, the FALCON-Unzip tool finds heterozygous single nucleotide polymorphisms and short indels in the haplotype-fused contigs and uses the phasing information within individual reads to group the reads into phase blocks. The read groups are reassembled and integrated with the remaining haplotype-fused contigs resulting in primary contigs and associated haplotigs.

Using error-corrected reads FALCON generated 4,774 sequences for a total of 2.91Gb of sequence with a contig N50 of 4.87Mb. This is comparable to the initial NA19240 assembly produced before integration with BioNano and alignment to GRCh38 to create chromosomal scaffolds. After running FALCON-Unzip, there were 2,230 Primary contig sequences and 9,916 haplotig sequences (Table S4). The Primary contig N50 is 4.95Mb while the haplotig contig N50 is considerably smaller at 440kb.

We aligned the haplotigs to the Hs-NA19240-1.0 using *nucmer* and looked for places where the haplotig had high-quality alignments on two or more chromosome scaffolds and/or unplaced scaffolds. We identified 245 unique haplotigs characterized by this split alignment pattern, and over 90% of these loci overlap segmental duplications. These represent two types of data: first, these represent places where the assembly algorithm collapsed the two haplotypes creating a mosaic representation of both haplotypes in a linear scaffold. Second, these represent places where segmental duplication and variation was too great between the two haplotypes, and during scaffolding to chromosomes, one haplotype was split into an unlocalized scaffold (Figure 4). In these cases, the haplotig analysis allows us to incorporate the unlocalized scaffold into the alternate haplotype representation further improving quality of the assembly. In Figure 4, an unlocalized scaffold fills a gap on chromosome 8 of the alternative haplotype in the *ASAP1/ADCY8* region. *ASAP1* encodes the ADP-ribosylation factor GTPase-activating protein, and decreased expression of *ASAP1* has been linked to predisposition to M. tuberculosis infection (Curtis et al. 2015). *ADCY8* is linked to short term episodic memory performance (de Quervain and Papassotiropoulos 2006). These data suggest that diploid assembly/phasing is valuable to complex genome assemblies as the method allows many sequences that were binned into unlocalized scaffolds during linear compression to haploid representation to be properly placed in their chromosomal context.

**Figure 4.**
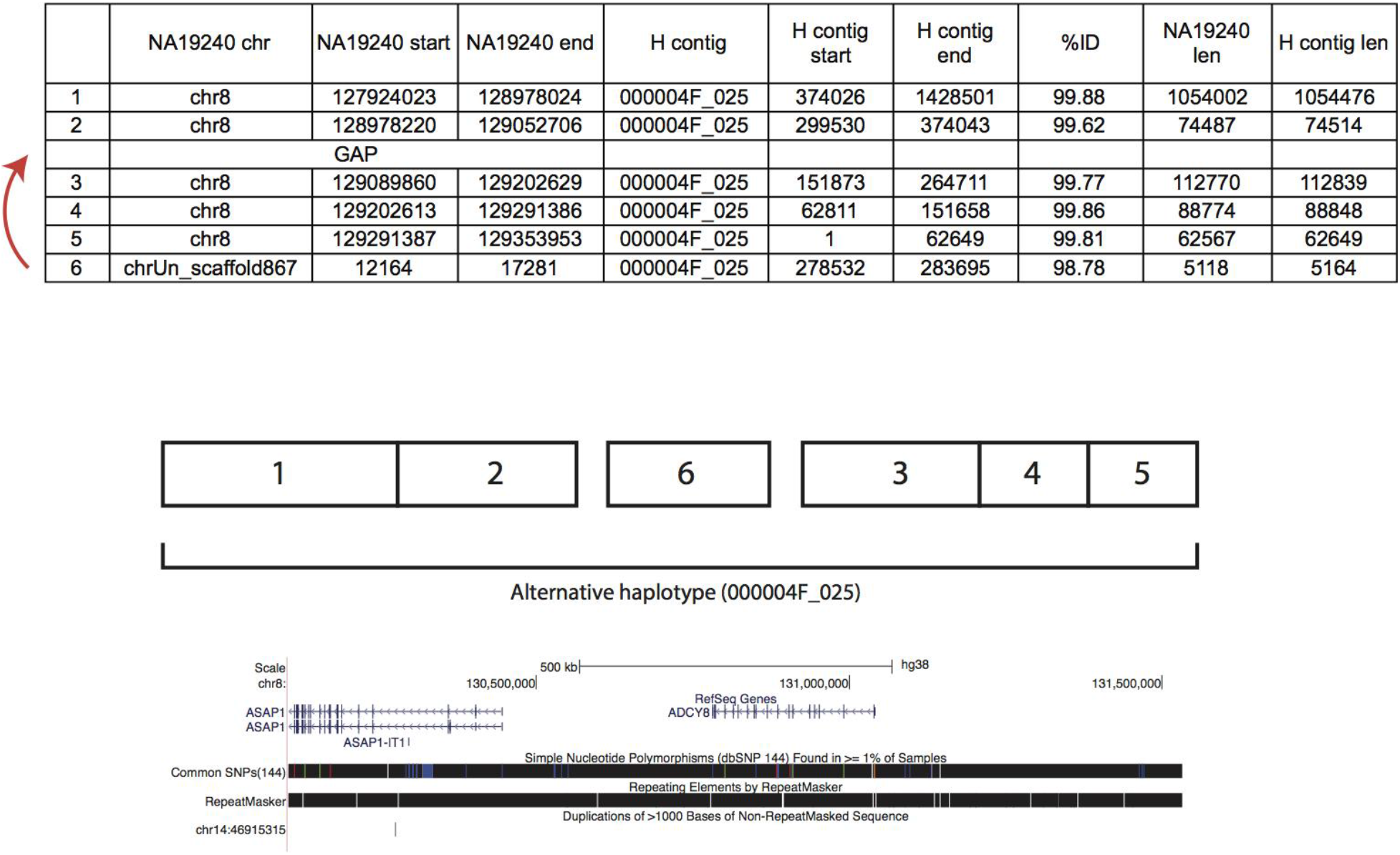
FALCONUnzip places unlocalized scaffolds in alternate haplotypes. The *nucmer* alignments of Hs-NA19240-1.0 to H contig 000004F_025 show that although most of the alignments are from chromosome 8, there is one alignment of an unlocalized scaffold (chrUn_scaffold867) as well as a gap between alignments #2 and #3. Placing the unlocalized scaffold (alignment #6) in the gap closes this gap and completes the alternative haplotype over the *ASAP1/ADCY8* region of chromosome 8 in Hs-NA19240-1.0.

## DISCUSSION

The general approach for genome analysis has been to utilize multiple resources to develop a haploid reference genome assembly which is used to support population level genome analysis. Reference-based analysis strategies allow scientists to align sequencing reads and call variants relative to the reference and have been the cornerstone of genome analysis for over a decade. While we have learned much from this approach, it is clear we do not fully characterize samples using this method. However, *de novo* genome assembly remains a significant challenge despite increase in throughput and decrease of sequence cost over the past decade. At the time of publication, the ‘finished’ human genome (NCBI35; GCF_000001405.11) contained 288 assembly gaps, and many of them were determined to be in regions containing structurally variant alleles. Additionally since the reference genome is assembled from clone sequence from many donors, it suffers from linear compression of haplotypes into a single-haplotype mosaic. For example, in the *MAPT* region, the allele represented in assembly versions prior to GRCh37 was not likely present in any individual human as it had been constructed by mixing the direct and inverted haplotypes present in the RP11 donor. BAC clone tiling paths from a single-haplotype source (CHM1) were used to resolve this region in the reference (Itsara et al. 2012), and alternative haplotypes were characterized from all continental populations (Steinberg et al. 2012). Accurately representing loci such as this allows researchers to query regions that were previously impossible to analyze due to the limitations of the available sequencing technologies, complex genome architecture, missing sequences, and various errors in the assembly or the underlying sequence data.

In this paper we describe our efforts to sequence and finish the genome of an African individual that is highly contiguous, approaching the quality of a reference assembly. We used a combination of sequencing and mapping technologies in addition to novel bioinformatic algorithms to construct this assembly and characterize regions of high structural diversity. Finally we use novel approaches to develop a haplotype-resolved assembly that more accurately represents a diploid genome.

This *de novo* assembly is the first of five assemblies from Africa, Europe, China, and Puerto Rico that we are working on towards the goal of adding more diversity to the reference genome. We characterize these assemblies as “gold genomes”, or highly contiguous representations of genomes with haplotype resolution of critical regions characterized by complex genetic architecture. These regions include beta-defensin, cytochrome P450 loci, and other segmentally duplicated loci with biomedical relevance. In comparison, a “platinum genome” is a contiguous, haplotype-resolved representation of the entire genome. To achieve these goals, we combine multiple sequencing technologies and analytic approaches such as PacBio SMRT sequencing, BioNano genomic maps, Illumina sequencing, 10x Genomics linked reads, and BAC hybridization and sequencing. We use all of these resources given that none of these approaches alone can fully resolve every genomic feature and/or region. For example, although PacBio provides long reads it still cannot traverse through many segmental duplications (Chaisson et al. 2015a)(Gordon et al. 2016); for this reason, we utilize BAC tiling paths across these loci. Although labor intensive and more expensive, the BAC library construction, hybridization, and sequencing is critical to our goal of haplotype resolution of complex regions of biomedical importance. Additionally, haplotype-resolved data is vital to improving our gold assemblies. For example, the linear compression of Hs-NA19240-1.0 to a haploid consensus led to an excess of frameshifted RefSeq transcripts compared to other WGS assemblies. Using the 10x Genomics direct diploid assembly we are able to distinguish between those transcripts that are likely due to error and those that may have biological context. The FALCON-Unzip algorithm also identified loci where the Hs-NA19240-1.0 assembly collapsed relevant information leading to a consensus primary contig and unplaced scaffolds. These unplaced scaffolds could then be recovered and placed in their genomic context as an alternative haplotype. We plan to integrate all of these data types (BAC, 10x Genomics, and FALCON-Unzip) into the next iteration of Hs-NA19240-1.0. For more in depth diploid analyses of NA19240, and an updated version of the Supernova™ assembly see Jaffe et al. (in preparation), a complementary manuscript to ours.

Incorporating standing variation into reference-based analyses is critical as it greatly improves accuracy in regions characterized by highly identical segmental duplications. Dilthey et al. (Dilthey et al. 2015) represent known variation in a population reference graph (PRG), a directed acyclic graphical model for genetic information that is generated using multiple sequence alignment of reference sequences and incorporates single nucleotide and short indel variation to all paths with matching sequence at that particular position. The authors then infer the diploid path through the PRG that most closely matches two haplotypes using a hidden Markov model. They use the MHC locus as an example and show that compared to classic SNP chip analyses and short-read and long-read mapping the PRG approach improves the accuracy of genotype inference particularly in regions of lower coverage sequencing and higher paralogy. This approach demonstrates the power of incorporating diversity to improve genome inference over traditional reference-based genotyping approaches.

One major challenge facing the genomics community is that although the current human reference includes representations for population diversity in the form of alternate loci scaffolds, most existing sequence analysis tools still expect a haploid assembly and cannot accurately handle a multi-allelic reference (Church et al. 2015). Given the level of diversity within the human populations, an optimal reference assembly could be represented by a graph in which shared sequences are depicted by nodes while population- and individual-specific sequences are depicted by edges and branch points in the graph. This solution allows for the representation of all available sequence in a contextual way, but also requires development of new analysis, annotation and display tools. Recently, a number of groups have proposed graph structures for representing the reference assembly (Church et al. 2011; Paten et al. 2014) and more specifically, Iqbal et al. (Iqbal et al. 2012) proposed the de Bruijn graph for pan genome analysis. Further, Marcus et al. (Marcus et al. 2014) developed a novel algorithm, splitMEM, for constructing a compressed de Bruijn graph (where the nodes and edges are compressed whenever the path between nodes is non-branching) from a generalized suffix tree of input genomes. This algorithm decomposes the Minimal Exact Matches (MEMs) from the suffix tree and extracts overlapping components to compute the nodes.

A number of novel approaches have recently emerged to utilize a graph reference. For example, BWA-MEM will map short reads to a primary assembly and the alternative scaffolds (Li 2013). The VG project has developed tools for creating and aligning to a graph and then calls variants relative to the graph (<u>https://github.com/vgteam/vg</u>). Finally, Rand et al. (Rand et al. 2016) describe an offset-based coordinate system that represents genomic intervals that includes all paths covered by the interval and the start and end coordinates allowing the full utilization of a graph reference assembly. In sum, the development of these novel tools complements our current efforts to add diversity to the reference assembly via alternative loci.

## MATERIALS AND METHODS

DNA for sequencing was purchased from Coriell Institute, and the cell line, GM19240, used to create the BioNano map, was also purchased from Coriell Institute. Cells were grown in AmnioMax C-100 Complete Medium (ThermoFisher Scientific) under 5% CO_2_ at 37°C. Methods for PacBio sequencing are in Supplementary Methods and Table S5. The initial assembly for NA19240 was performed at DNANexus using FALCON and Quiver. The parameters for FALCON and Quiver as well as the DISCOVARDeNovo and DISCOVARDeNovo-SSPACE assemblies are in Supplementary Methods.

### BioNano

See Supplementary Methods and Table S6 for detailed BioNano library preparation methods. Hs-NA19240-1.0 was nicked in silico with BspQI to produce a cmap file, which reports the start and end coordinates and the placement of labels for each contig. BioNano Genomics software tools were then used to align BioNano genome maps to NA19240 PacBio unitigs and produce ultra-long Hybrid Scaffolds. This software also identifies and reports “conflicts”, defined as genome maps and PacBio contigs that align to one another with a significant number of unaligned labels outside an aligned region, often implying a misassembly or chimera in either the BioNano genome map or the sequence assembly. These conflicting junctions were manually reviewed to identify potential breaks in the PacBio unitigs.

### Reference-Guided Chromosome Assembly

The ranked alignments generated from an NCBI assembly-assembly alignment of WGS assembly LKPB01000000 and GRCh38 were used as input for the assembly of chromosome sequences for Hs-NA19240-1.0. For each chromosome, we generated a tiling path along which the order and orientation of NA19240 contigs reflects their alignment to the corresponding GRCh38 chromosome. See Supplementary Methods for filtering criteria. We used the finalized TPFs and pair-wise alignments to define switch points and create an AGP file for each chromosome.

### BAC Library

The VMRC64 BAC library was constructed using the methods in Smith et al. (Smith et al. 2010). High molecular weight DNA was isolated and embedded into agarose blocks, then partially digested with EcoRI and subcloned into the pCC1BAC vector (Epicentre) to create >150 kbp insert libraries. Clones were plated on LB/1.5% agar plates supplemented with 12.5 ug/mL chloramphenicol, 120 ug/mL X-Gal and 0.1mM IPTG, picked into 384-well microtiter plates, then transferred to high density nylon filters for library screening. Clones were hybridized using the overgo protocol from BACPAC (<u>https://bacpac.chori.org/overgohyb.htm</u>), and then sequenced using either PacBio or Illumina MiSeq technologies. The sequenced clones from this library are accessioned in GenBank: <u>https://www.ncbi.nlm.nih.gov/clone/library/genomic/37646175/</u>.

### Fosmid Library

ABC10 fosmid clone placements were performed and evaluated as described in (Steinberg et al. 2014b; Schneider et al. 2013b). On the GCA_001524155.1 assembly, the average insert length = 39,448.1 and the standard deviation = 4,570.7.

### Assembly-assembly alignment and RefSeq transcript alignment

Assemblies were also aligned using software version 1.7 of the NCBI pipeline as described in the Methods section of (Steinberg et al. 2014a). The alignment and alignment reports are available from the NCBI Remap FTP site: <u>ftp://ftp.ncbi.nlm.nih.gov/pub/remap/Homosapiens/1.7/</u> (Kitts et al. 2015). We evaluated chromosome-level collapse and expansion in these alignments with <u>https://github.com/deannachurch/assemblyalignment/</u>. Alignments were performed and analyzed as described in the Supplementary Methods of (Shi et al. 2016). See Supplementary Methods for detailed parameters on NCBI assembly-assembly alignment.

### FALCONUnzip and 10x Genomics

Parameters used for FALCONUnzip can be found in Supplementary Methods. Detailed methods for the NA19240_10x_dev assembly can be found in Jaffe et al. (in preparation).

## DATA ACCESS

Hs-NA19240-1.0 in GenBank: GCA_001524155.1

PacBio sequence reads: SRX1094388, SRX1096798, SRX1094289, SRX1094374, SRX1093654, SRX1093555, SRX1093000

Illumina PCR free sequence reads: SRX1098166

3Kb mate-pair data (SRX1098164, SRX1098165)
550bp insert paired-end reads (SRX1098162, SRX1098163)

DISCOVARDeNovo and DISCOVARDeNovo-SSPACE assembly in GenBank: submitted awaiting accessions

NA19240_10x_dev assembly available at https://www.dropbox.com/sh/8gsdycmqk04ei7s/AADZJgJtcK_mw16CYpoSpJhca?dl=0

## ACKNOWLEDGEMENTS AND DISCLOSURES

We would like to thank Aye Wollam and Susie Rock for their work on clone-based finishing at McDonnell Genome Institute. We would also like to thank Jason Chin for generating the FALCON-Unzip data. Many thanks to Tonia Brown for careful reading and editing of the manuscript. This work was supported, in part, by grants from the U.S. National Institutes of Health (NIH grant 5U41HG007635 to RKW and NIH grant HG002385 to EEE). EEE is an investigator of the Howard Hughes Medical Institute. EEE is on the scientific advisory board (SAB) of DNAnexus, Inc., is a consultant for Kunming University of Science and Technology (KUST) as part of the 1000 China Talent Program, and was an SAB member of Pacific Biosciences, Inc. (2009–2013). DMC is an employee of 10X Genomics.

## REFERENCES

1000 Genomes Project Consortium, Auton A, Brooks LD, Durbin RM, Garrison EP, Kang HM, Korbel JO, Marchini JL, McCarthy S, McVean GA, et al. 2015. A global reference for human genetic variation. Nature 526: 68–74.

Antonacci F, Dennis MY, Huddleston J, Sudmant PH, Steinberg KM, Rosenfeld JA, Miroballo M, Graves TA, Vives L, Malig M, et al. 2014. Palindromic GOLGA8 core duplicons promote chromosome 15q13.3 microdeletion and evolutionary instability. Nat Genet 46: 1293–1302.

Antonacci F, Kidd JM, Marques-Bonet T, Ventura M, Siswara P, Jiang Z, Eichler EE. 2009. Characterization of six human disease-associated inversion polymorphisms. Hum Mol Genet 18: 2555–2566.

Bailey JA, Gu Z, Clark RA, Reinert K, Samonte RV, Schwartz S, Adams MD, Myers EW, Li PW, Eichler EE. 2002. Recent segmental duplications in the human genome. Science 297: 1003–1007.

Boetzer M, Henkel CV, Jansen HJ, Butler D, Pirovano W. 2011. Scaffolding pre-assembled contigs using SSPACE. Bioinformatics 27: 578–579.

Chaisson MJP, Huddleston J, Dennis MY, Sudmant PH, Malig M, Hormozdiari F, Antonacci F, Surti U, Sandstrom R, Boitano M, et al. 2015a. Resolving the complexity of the human genome using single-molecule sequencing. Nature 517: 608–611.

Chaisson MJP, Wilson RK, Eichler EE. 2015b. Genetic variation and the de novo assembly of human genomes. Nat Rev Genet 16: 627–640.

Chin C-S, Alexander DH, Marks P, Klammer AA, Drake J, Heiner C, Clum A, Copeland A, Huddleston J, Eichler EE, et al. 2013. Nonhybrid, finished microbial genome assemblies from long-read SMRT sequencing data. Nat Methods 10: 563–569.

Chin C-S, Peluso P, Sedlazeck FJ, Nattestad M, Concepcion GT, Clum A, Dunn C, O’Malley R, Figueroa-Balderas R, Morales-Cruz A, et al. 2016. Phased Diploid Genome Assembly with Single Molecule Real-Time Sequencing. http://biorxiv.org/lookup/doi/10.1101/056887.

Church DM, Schneider VA, Graves T, Auger K, Cunningham F, Bouk N, Chen H-C, Agarwala R, McLaren WM, Ritchie GRS, et al. 2011. Modernizing reference genome assemblies. PLoSBiol 9: e1001091.

Church DM, Schneider VA, Steinberg KM, Schatz MC, Quinlan AR, Chin C-S, Kitts PA, Aken B, Marth GT, Hoffman MM, et al. 2015. Extending reference assembly models. Genome Biol 16: 13.

Curtis J, Luo Y, Zenner HL, Cuchet-Loureno D, Wu C, Lo K, Maes M, Alisaac A, Stebbings E, Liu JZ, et al. 2015. Susceptibility to tuberculosis is associated with variants in the ASAP1 gene encoding a regulator of dendritic cell migration. Nat Genet 47: 523–527.

Dennis MY, Nuttle X, Sudmant PH, Antonacci F, Graves TA, Nefedov M, Rosenfeld JA, Sajjadian S, Malig M, Kotkiewicz H, et al. 2012. Evolution of human-specific neural SRGAP2 genes by incomplete segmental duplication. Cell 149: 912–922.

de Quervain DJ-F, Papassotiropoulos A. 2006. Identification of a genetic cluster influencing memory performance and hippocampal activity in humans. Proc Natl Acad Sci U S A 103: 4270–4274.

Dilthey A, Cox C, Iqbal Z, Nelson MR, McVean G. 2015. Improved genome inference in the MHC using a population reference graph. Nat Genet 47: 682–688.

Florea L, Souvorov A, Kalbfleisch TS, Salzberg SL. 2011. Genome assembly has a major impact on gene content: a comparison of annotation in two Bos taurus assemblies. PLoS One 6: e21400.

Gnerre S, Maccallum I, Przybylski D, Ribeiro FJ, Burton JN, Walker BJ, Sharpe T, Hall G, Shea TP, Sykes S, et al. 2011. High-quality draft assemblies of mammalian genomes from massively parallel sequence data. Proc Natl Acad Sci U S A 108: 1513–1518.

Gordon D, Huddleston J, Chaisson MJP, Hill CM, Kronenberg ZN, Munson KM, Malig M, Raja A, Fiddes I, Hillier LW, et al. 2016. Long-read sequence assembly of the gorilla genome. Science 352: aae0344.

Iqbal Z, Caccamo M, Turner I, Flicek P, McVean G. 2012. De novo assembly and genotyping of variants using colored de Bruijn graphs. Nat Genet 44: 226–232.

Itsara A, Vissers LELM, Steinberg KM, Meyer KJ, Zody MC, Koolen DA, de Ligt J, Cuppen E, Baker C, Lee C, et al. 2012. Resolving the breakpoints of the 17q21.31 microdeletion syndrome with next-generation sequencing. Am J Hum Genet 90: 599–613.

Kitts PA, Church DM, Thibaud-Nissen F, Choi J, Hem V, Sapojnikov V, Smith RG, Tatusova T, Xiang C, Zherikov A, et al. 2015. Assembly: a resource for assembled genomes at NCBI. Nucleic Acids Res. http://dx.doi.org/10.1093/nar/gkv1226.

Kurtz S, Phillippy A, Delcher AL, Smoot M, Shumway M, Antonescu C, Salzberg SL. 2004. Versatile and open software for comparing large genomes. Genome Biol 5: R12.

Li H. 2013. Aligning sequence reads, clone sequences and assembly contigs with BWA-MEM. arXiv [q-bioGN]. http://arxiv.org/abs/1303.3997.

Marcus S, Lee H, Schatz MC. 2014. SplitMEM: a graphical algorithm for pan-genome analysis with suffix skips. Bioinformatics 30: 3476–3483.

Mostovoy Y, Levy-Sakin M, Lam J, Lam ET, Hastie AR, Marks P, Lee J, Chu C, Lin C, Dzakula Z, et al. 2016. A hybrid approach for de novo human genome sequence assembly and phasing. Nat Methods 13: 587–590.

Nattestad M, Schatz MC. 2016. Assemblytics: a web analytics tool for the detection of assembly-based variants. http://biorxiv.org/lookup/doi/10.1101/044925.

Paten B, Novak A, Haussler D. 2014. Mapping to a Reference Genome Structure. arXiv [q-bioGN]. http://arxiv.org/abs/1404.5010.

Paulino D, Warren RL, Vandervalk BP, Raymond A, Jackman SD, Birol I. 2015. Sealer: a scalable gap-closing application for finishing draft genomes. BMC Bioinformatics 16: 230.

Pendleton M, Sebra R, Pang AWC, Ummat A, Franzen O, Rausch T, Stütz AM, Stedman W, Anantharaman T, Hastie A, et al. 2015. Assembly and diploid architecture of an individual human genome via single-molecule technologies. Nat Methods 12: 780–786.

Rand KD, Grytten I, Nederbragt A, Storvik GO, Glad IK, Sandve GK. 2016. Coordinates and Intervals in Graph-based Reference Genomes. http://biorxiv.org/lookup/doi/10.1101/063206.

Schneider VA, Chen H-C, Clausen C, Meric PA, Zhou Z, Bouk N, Husain N, Maglott DR, Church DM. 2013a. Clone DB: an integrated NCBI resource for clone-associated data. Nucleic Acids Res 41: D1070–8.

Schneider VA, Chen HC, Clausen C, Meric PA, Zhou Z, Bouk N, Husain N, Maglott DR, Church DM. 2013b. Clone DB: an integrated NCBI resource for clone-associated data. Nucleic Acids Res 41: D1070–8.

She X, Jiang Z, Clark RA, Liu G, Cheng Z, Tuzun E, Church DM, Sutton G, Halpern AL, Eichler EE. 2004. Shotgun sequence assembly and recent segmental duplications within the human genome. Nature 431: 927–930.

Shi L, Guo Y, Dong C, Huddleston J, Yang H, Han X, Fu A, Li Q, Li N, Gong S, et al. 2016. Long-read sequencing and de novo assembly of a Chinese genome. Nat Commun 7: 12065.

Smit AFA, Hubley R, Green P. 1996. RepeatMasker. Published on the web at http://www.repeatmasker.org.

Smith JJ, Stuart AB, Sauka-Spengler T, Clifton SW, Amemiya CT. 2010. Development and analysis of a germline BAC resource for the sea lamprey, a vertebrate that undergoes substantial chromatin diminution. Chromosoma 119: 381–389.

Steinberg KM, Antonacci F, Sudmant PH, Kidd JM, Campbell CD, Vives L, Malig M, Scheinfeldt L, Beggs W, Ibrahim M, et al. 2012. Structural diversity and African origin of the 17q21. 31 inversion polymorphism. Nat Genet 44: 872–880.

Steinberg KM, Schneider VA, Graves-Lindsay TA, Fulton RS, Agarwala R, Huddleston J, Shiryev SA, Morgulis A, Surti U, Warren WC, et al. 2014a. Single haplotype assembly of the human genome from a hydatidiform mole. Genome Res 24: 2066–2076.

Watson CT, Steinberg KM, Graves TA, Warren RL, Malig M, Schein J, Wilson RK, Holt RA, Eichler EE, Breden F. 2015. Sequencing of the human IG light chain loci from a hydatidiform mole BAC library reveals locus-specific signatures of genetic diversity. Genes Immun 16: 24–34.

Watson CT, Steinberg KM, Huddleston J, Warren RL, Malig M, Schein J, Willsey AJ, Joy JB, Scott JK, Graves TA, et al. 2013. Complete Haplotype Sequence of the Human Immunoglobulin Heavy-Chain Variable, Diversity, and Joining Genes and Characterization of Allelic and Copy-Number Variation. Am J Hum Genet 92: 530–546.

Weisenfeld NI, Yin S, Sharpe T, Lau B, Hegarty R, Holmes L, Sogoloff B, Tabbaa D, Williams L, Russ C, et al. 2014. Comprehensive variation discovery in single human genomes. Nat Genet 46: 1350–1355.

Zheng GXY, Lau BT, Schnall-Levin M, Jarosz M, Bell JM, Hindson CM, Kyriazopoulou-Panagiotopoulou S, Masquelier DA, Merrill L, Terry JM, et al. 2016. Haplotyping germline and cancer genomes with high-throughput linked-read sequencing. Nat Biotechnol 34: 303–311.

